# Sketching Algorithms for Genomic Data Analysis and Querying in a Secure Enclave

**DOI:** 10.1101/468355

**Authors:** Can Kockan, Kaiyuan Zhu, Natnatee Dokmai, Nikolai Karpov, M. Oguzhan Kulekci, David P. Woodruff, S. Cenk Sahinalp

## Abstract

Current practices in collaborative genomic data analysis (e.g. PCAWG [1]) necessitate all involved parties to exchange individual patient data and perform all analysis locally, or use a trusted server for maintaining all data to perform analysis in a single site (e.g. the Cancer Genome Collaboratory). Since both approaches involve sharing genomic sequence data - which is typically not feasible due to privacy issues, collaborative data analysis remains to be a rarity in genomic medicine.

In order to facilitate efficient and effective collaborative or remote genomic computation we introduce `SkSES` (Sketching algorithms for Secure Enclave based genomic data analysiS), a computational framework for performing data analysis and querying on multiple, individually encrypted genomes from several institutions in an untrusted cloud environment. Unlike other techniques for secure/privacy preserving genomic data analysis, which typically rely on sophisticated cryptographic techniques with prohibitively large computational overheads, `SkSES` utilizes the secure enclaves supported by current generation microprocessor architectures such as Intel’s SGX. The key conceptual contribution of `SkSES` is its use of *sketching* data structures that can fit in the limited memory available in a secure enclave.

While streaming/sketching algorithms have been developed for many applications in computer science, their feasibility in genomics has remained largely unexplored. On the other hand, even though privacy and security issues are becoming critical in genomic medicine, available cryptographic techniques based on, e.g. homomorphic encryption or garbled circuits, fail to address the performance demands of this rapidly growing field. The alternative offered by Intel’s SGX, a combination of hardware and software solutions for secure data analysis, is severely limited by the relatively small size of a secure enclave, a private region of the memory protected from other processes. `SkSES` addresses this limitation through the use of sketching data structures to support efficient secure and privacy preserving SNP analysis across individually encrypted VCF files from multiple institutions. In particular `SkSES` provides the users the ability to query for the “*k* most significant SNPs” among any set of user specified SNPs and any value of *k* - even when the total number of SNPs to be maintained is far beyond the memory capacity of the secure enclave.

**Results:** We tested `SkSES` on the complete iDASH-2017 competition data set comprised of 1000 case and 1000 control samples related to an unknown phenotype. `SkSES` was able to identify the top SNPs with respect to the *χ*^2^ statistic, among any user specified subset of SNPs across this data set of 2000 individually encrypted complete human genomes quickly and accurately - demonstrating the feasibility of secure and privacy preserving computation for genomic medicine via Intel’s SGX.

**Availability:** https://github.com/ndokmai/sgx-genome-variants-search

## 1 Introduction

As genome sequencing technologies get faster and cheaper, world-wide growth rate for genomic data has surpassed Moore’s law - which indicates that the size of memory that can be fitted in a chip doubles every 18 months. As a result, computational methods that reduce memory usage, e.g., by representing genomic data more compactly, or making inference on the fly by processing genomic data in an on-line manner are of high demand. Among these approaches, lossless compression methods on raw, mapped or indexed data [2, 3, 4, 5, 6, 7] have been highly successful; for example, recent standardization efforts by MPEG-G [8, 9] or GA4GH [10]), especially in the context of raw (FASTQ) and mapped (SAM/BAM) read collections based on current generation compression methods, have demonstrated that it is possible to reduce the size of genomic data by an order of magnitude. Similarly, on-line genomic data processing methods (e.g. eXpress [11], Sailfish [12] and Salmon [13]) have demonstrated how to make inference on large scale sequence data by treating it as a “stream”, typically using smaller memory than conventional techniques. Interest in the on-line treatment of biomolecular sequence data has also lead to a standardization effort by GA4GH, with the aim of improving the ability to share and process large scale genomic data as a “stream” [14].

The first algorithms for processing sequential data through the use of limited memory (too small to store all data) were devised in late 1970s [15]. However, the first streaming algorithms, which aim to make inference on a data collection in a single pass while using sublinear memory, were described much later [16]. The study of data streams eventually gave rise to “sketching” algorithms [17, 18] which summarize certain characteristics of input data in the form of a *sketch*; such algorithms differ from the on-line methods mentioned above because they process an entire data set in a limited memory setting before making any inference. Sketching algorithms also differ from streaming methods in the way that they can access data more than once. As such, they are particularly useful for multi-level storage systems where the bulk of data is stored in the slow, possibly external storage while the fast working memory maintains a sketch of the data on which inferences are made.

As genomic data grow in size and importance, so does the need to perform its analysis in a privacy preserving and secure manner. Genomic data carry information about, e.g. predisposition to disease [19], not only of the people from whom such data is collected, but also close relatives [20]. Once made public, genomic data (just like other biometric data) becomes irrevocable and may imply future privacy risks with further discoveries in human genetics [21]. Collaborative analysis of genomic data involving multiple parties (research institutions, government organizations, countries) is further complicated by policies that may limit or fully prohibit sharing of genomic data from individuals.

There are a number of available cryptographic techniques [22, 23, 24, 25, 26, 27, 28] that have been proposed to address some of the above concerns. Unfortunately, due to their high computational overhead, none of these techniques can scale up to address the performance needs of real-life genomic analysis [29, 30, 31, 32]. For example, the use of homomorphic encryption (HME) [33] can help data owners securely outsource genomic data analysis; unfortunately HME is slow and memory intensive thus can only work for small datasets. Similarly, secure multiparty computation (SMC) [34] offers an interesting platform for secure collaboration; unfortunately it comes with even higher computational overhead. Garbled circuit-based methods (e.g. [35]) require sophisticated circuit design and optimization for each specific task, resulting in limited/no flexibility. Finally secret sharing-based secure genome-wide association methods [24] offer better performance than HME and garbled circuit-based methods, yet they still have significant computation overhead when compared to the same analysis on plaintext. As summarized above, existing software based solutions to secure genomic analysis rely on heavyweight cryptographic techniques. On the other hand, Intel’s Software Guard Extensions (SGX) [36], a combined hardware and software platform supported by all current generation Intel processor architectures, offer users sensitive data analysis within a protected enclave. Solutions based on SGX are not expected to introduce significant computational overhead or restrictions on basic operations common to software-based techniques such as the garbled circuit-based FlexSC framework [35] or homomorphic encryption based HElib framework [33], and thus is likely to make secure genome-scale data analysis feasible.

The type of analysis we focus on in this paper is Genome-Wide Association Studies (GWAS), a collection of statistical methods that aim to associate certain genome alterations and rearrangements with certain conditions [37]. Perhaps the most commonly used statistic to associate, e.g. Single Nucleotide Polymorphisms (SNPs) with certain conditions is the *χ*^2^ statistic [38]. As per many others, the *χ*^2^ statistic is highly dependent on the population size; thus larger and more diverse cohorts are typically preferred. Bringing together such a cohort, especially for rare diseases, may necessitate multiple institutions to collaborate and share data to collectively calculate these statistics for developing an accurate model.

In this paper, we present `SkSES` (Sketching algorithms for Secure Enclave based genomic data analysiS - pronounced “success”), a computational framework to perform secure collaborative genomic analysis, in particular GWAS, and on-line querying, on an untrusted cloud platform with minimal performance overhead, through the use of Intel’s SGX architectures (as well as other similar platforms that are under development). Since the secure enclave supported by this platform has significant memory limitations and a large overhead for loading/unloading data, efficient compaction of both the genomic data to be processed and the data structure to perform inference (for this case computing the statistic of interest) are of utmost importance.

For compacting the SNP data provided in Variant Call Format (VCF), we present a simple compression scheme to filter out non-essential components of an individual VCF file and encode essential components efficiently - which reduces the storage and communication needs but also speeds up the encryption and decryption procedures within the framework. For compacting the data structure to maintain SNP frequencies, for the purpose of computing the statistic of interest and identifying the most significant SNPs, we present a “sketching” approach through which we can identify the top *k* SNPs (*k* is any reasonably small value which can be predefined or can be defined during querying) among a user specified subset of SNPs across case/control samples. As we will show, our new sketching approach is not only more efficient than conventional data storage techniques - significantly improving on our iDASH-2017 (http://www.humangenomeprivacy.org/2017/) runner-up software for secure GWAS, but also allows the users to pose on-line queries, which are not possible by standard data storage approaches within the constraints of the secure enclave.

## 2 Methods

### 2.1 Overview of the SGX Platform

SGX is a security extension of Intel processor architecture. By using SGX, privileged modules like operating system (OS), virtual machine (VM) scheduler etc. are isolated from private codes and secret data through hardware protection. More specifically, instead of quarantining malicious parts within the running system Intel SGX seals private codes, sensitive data and other selected secrets into a CPU secure computation unit called “enclave”. The access of secrets within enclaves are restricted by the hardware supported access control.

A typical SGX based application consists of (one or more) data owner(s), an untrusted cloud service provider (CSP), and the secure enclave. First, the data owner establishes a secure channel with the enclave hosted by an untrusted CSP through the remote attestation process. Then, the data owner can securely upload data to the CSP (data provisioning). In SGX, all decrypted secrets can only be accessed by the authorized codes, which also lie inside the enclave. A hardware supported access control proxy guarantees the code and data cannot be accessed or modified by any software outside the secure enclave.

### 2.2 SkSES Setup

`SkSES` set up involves individual genomic data in the form of VCF files from one or more parties who would like to perform statistical tests on the entire data set. We report on results with VCF files specifying SNPs from chromosome 1 only (as per iDASH-2017 competition) as well as the entire human genome data. Each VCF file is marked as either case or control and is individually filtered, compressed and encrypted as will be described, and is uploaded to an untrusted CSP. The specific analysis/querying offered by `SkSES` to the users is, given a user specified set of SNPs (which can be the entire set of SNPs in the human genome or a subset) and an integer *k*, what are the most significant *k* SNPs with respect to the commonly employed *χ*^2^ statistic within this case/control cohort? `SkSES` achieves this by first processing the entire data set to establish a “sketch” within the enclave. On a given query, `SkSES` identifies (a super-set of) potentially significant SNPs and re-accesses the relevant portions of the VCF files to identify the most significant *k* SNPs.

Note that the setup we consider here is a more general version of that for the iDASH-2017 competition in the sense that we allow the user to pose multiple on-line queries with specific value of *k* and the subset of SNPs among which the top *k* is identified by `SkSES` based on the *χ*^2^ statistic. ^1^

### 2.3 Filtering and Compression of VCF Files for GWAS

Mapped and processed genomic data with variant calls are typically stored as VCF files where each line contains information about a single variant. For each SNP, a VCF file includes “ID”, “TYPE”, “CHROM”, “POS”, “REF”, “ALT”, “QUAL”, and “FILTER” columns (and possibly more). For the purposes of GWAS (in particular the *χ*^2^ statistic) only “ID” and “TYPE” columns are mandatory: “QUAL” and “FILTER” parameters would be identical across all individuals participating in such analysis; “CHROM”, “POS”, “REF”, and “ALT” column information can all be obtained via the “ID” column from external resources such as the dbSNP database. Considering that each VCF file needs to be encrypted, sent over some channel, loaded into the SGX enclave, decrypted, and finally processed, reducing the size of the data can improve the overall performance significantly. To further reduce the size of the data, one can trim the “(r/s)s” in front of a SNP “ID”, and represent the remaining part as a 4-byte integer. The entries can then be sorted by “TYPE”, so as to represent the zygosity status for SNPs only onceand then by “ID” to obtain integer sequence 〈*x*_1_, *x*_2_,…*x_k_*〉 for a particular “TYPE”, where *x_i_* < *x*_*i*+1_ for any *i*. This makes it possible to use delta (differential) encoding on the “ID” sequence 〈*x*_1_, *x*_2_,…*x_k_*〉 so that it could be represented 〈*x*_1_, *δ*_1_, *δ*_2_,…, *δ*_*k*–1_〉, where *δ_i_* = *x*_*i*+1_ – *x_i_*. Each *δ_i_* is encoded with its corresponding minimal binary representation (MBR) by using ⌊log(*x*_*i*+1_ – *x_i_*)⌋ bits, omitting the most significant bit (since it is always 1). Although the bit string obtained by concatenating (variable length) MBR values (namely *payload*) is not uniquely decodable, we keep a companion bit string (namely *disambiguation information*) representing code word lengths ⌊log *δ_i_*⌋ for each *i* - which are further compressed through static Huffman encoding.

In summary our format asks to filter out all fields that are not essential for GWAS (in particular the *χ*^2^ statistic) on the data, partition the IDs according to zygosity status, keep the integer part of the IDs, sort and delta encode them in two streams. Additionally the format includes a boolean value indicating the case/control status, and an integer indicating where the data stream changes zygosity.

### 2.4 Identifying the Top SNPs Based on the *χ*^2^ Statistic in the Secure Enclave

Once the filtering/compression steps have been performed, the parties can exchange data and messages required for the actual computation. The tasks are divided between the server, which refers to the cloud service provider (CSP) equipped with Intel SGX supported hardware, and the client(s), which refer to individual users (e.g. genome centers) that want to perform their collaborative analysis in a privacy preserving and secure manner. As per the Intel SGX architecture guidelines, the server begins by creating and initializing an SGX enclave and waits for service requests from the clients. Then, a client needs to initiate the remote attestation protocol to establish that the server is indeed recognized by Intel and is going to perform the desired computation. Once the remote attestation is complete and the parties have verified each other and their intentions, the server allocates sufficient resources to performs the computation.

For our application the majority of the SGX enclave memory is used to keep either the basic data structures – in the form of hash tables maintaining allele counts (which is described in detail in Section 2.5), or the space-efficient data structures that are employed when the SGX memory cannot store all SNP IDs (which is described in detail in Section 2.7). As the server receives the compressed and encrypted data from the clients, the data structures residing in the SGX enclave are updated. Once all incoming data has been processed, the standard *χ*^2^ statistic can be computed for each SNP entry within the hash table to determine the top-*k* SNPs with respect to the *χ*^2^ statistic (see Definition 1 below). For the case that the SNP IDs and their corresponding allele counts exceed memory limit, `SkSES` identifies some *l* SNPs (*l* > *k*) which include the top-*k* SNPs with high probability, and processes the data in another pass to filter out all but the most significant *k* SNPs. Finally, the results are sent back to the clients.

#### Definition 1.

Ranking SNPs with respect to *χ*^2^ statistic [38].

**Input**: A collection of *m* sparse vectors **v**_*j*_ from participating parties, each with *n* dimensions, s.t. **v**_*j*_ [*i*] ∈ {0, 1, 2} (*i* ∈ [*n*] and *j* ∈ [*m*] - where [*k*] indicates {1, 2, ⋯, *k*}), representing the genotype of individual *j* for a particular variant *i*. Typically **v**_*j*_ [*i*] = 0 - indicating that the variant *i* is not present in *j*; **v**_*j*_[*i*] = 1/2 respectively indicate that the variant is heterozygous/homozygous. Additionally each **v**_*j*_ has a class label which is either 1 (case) or –1 (control), denoted as sign(*j*) = ±1. The number of case-labeled and control-labeled vectors are denoted as *R*, and *S* = *m* – *R* respectively.

**Querying:** We consider two types of queries.

1. Return each of the *k* “dimensions” *i* for which **C**[*i*] is one of the largest *k* values among **C**[1: *n*] where 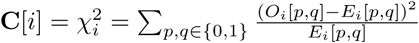 (see below for the definition of *O_i_* and *E_i_*) if ∀*p*, *q* ∈ {0, 1}, *E_i_*[*p*, *q*] ≠ 0 and 0 otherwise. In other words return rank(**C**)[1: *k*] where rank(**L**) is the sorted list of the *n* dimensions of a given a vector **L** ∈ ℝ^*n*^ in descending order.
2. For a user specified subset *T* ⊂ [*n*] (a subset of all possible SNP IDs), return the largest *k* dimensions in *T*, i.e. rank(**C**[*i* ∈ *T*])[1: *k*].

For each dimension *i* let the standard 2 × 3 table of genotype counts from the case and control groups be *G_i_*, where *r_ih_* = |{*j* | **v**_*j*_[*i*] = *h* ∧ sign(*j*) = 1}| and *s_ih_* = |{*j* | **v**_*j*_[*i*] = *h* ∧ sign(*j*) = –1}| respectfully represent the total count of each genotype *h* = 0, 1, 2 for a given *i*, from the case and control groups, and *n_ih_* = *r_ih_* + *s_ih_*.

| Genotype counts ($G_i$) | $\mathbf{v}_j[i] = 0$ | $\mathbf{v}_j[i] = 1$<br>(i.e. heterozygous) | $\mathbf{v}_j[i] = 2$<br>(i.e. homozygous) | Sum |
| --- | --- | --- | --- | --- |
| Cases ($\text{sign}(j) = 1$) | $r_{i0}$ | $r_{i1}$ | $r_{i2}$ | $R = r_{i0} + r_{i1} + r_{i2}$ |
| Controls ($\text{sign}(j) = -1$) | $s_{i0}$ | $s_{i1}$ | $s_{i2}$ | $S = s_{i0} + s_{i1} + s_{i2}$ |
| Sum | $n_{i0} = r_{i0} + s_{i0}$ | $n_{i1} = r_{i1} + s_{i1}$ | $n_{i2} = r_{i2} + s_{i2}$ | $m = R + S$ |

The following 2 × 2 tables define *O_i_* - the observed, and *E_i_* - the expected allele counts (obtained by summarizing *G_i_*), completing the definition of
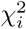.

| Observed ($O_i$) | Ref. | SNP | Sum |
| --- | --- | --- | --- |
| Cases | $2r_{i0} + r_{i1}$ | $2r_{i2} + r_{i1}$ | $2R$ |
| Controls | $2s_{i0} + s_{i1}$ | $2s_{i2} + s_{i1}$ | $2S$ |
| Sum | $2n_{i0} + n_{i1}$ | $2n_{i2} + n_{i1}$ | $2m$ |

| Expected ($E_i$) | Ref. | SNP | Sum |
| --- | --- | --- | --- |
| Cases | $R(2n_{i0} + n_{i1})/m$ | $R(2n_{i2} + n_{i1})/m$ | $2R$ |
| Controls | $S(2n_{i0} + n_{i1})/m$ | $S(2n_{i2} + n_{i1})/m$ | $2S$ |
| Sum | $2n_{i0} + n_{i1}$ | $2n_{i2} + n_{i1}$ | $2m$ |

### 2.5 Basic Data Structures in the Secure Enclave

Even though it offers a limited memory, Intel SGX secure enclave can still handle genomic data analysis in a limited scale (e.g. GWAS on a single human chromosome) through the use of standard (non-sketching) data structures. We have developed and tested several data structure options for computing the *χ*^2^ statistic on a limited set of SNPs from the human genome to provide a baseline performance - on SNPs from chromosome 1 (as per the iDASH-2017 competition). In particular we offer three hash table options to maintain for each variant (SNP ID) *i* the values *r*_*i*1_ + 2*r*_*i*2_ and *s*_*i*1_ + 2*s*_*i*2_) necessary for the calculation of
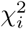 (see equation (1) in Section 2.6): (i) a standard open-address hash table, (ii) a chained hash table with move-to-front heuristic, and (iii) a Robin-Hood (open-address) hash table.

A chained hash table with move-to-front heuristic modifies the standard chained hash table by moving the last updated table entry to the front of the associated linked list (i.e. chain). Such a hash table performs especially well when the data is highly skewed (e.g. follows a distribution that is closer to Zipfian rather than uniform - we discuss the hypothetical distribution of SNP counts in Section 2.6). Its main drawback for our application is that the pointers lead to heap fragmentation, which could be critical for the execution within the SGX enclave. The alternative Robin-Hood hash table [40] modifies the standard open-address hash table in a way that each new entry is swapped with an existing entry if the “probe length” of the new entry is higher; since there are no deletions performed, we use linear probing.

### 2.6 Computing the top-*k* SNPs with respect to *χ*^2^ Statistic via Sketching

Maintaining all SNPs in the human genome and their respective allelic counts exceeds the memory limit of the secure enclave.^2^ In other words, the working memory of the SGX enclave has *n*′ = *o*(*n*) words (i.e. *o*(*n*(log *m* + log *n*)) bits). A straightforward solution in this case is to partition the *n* dimensions (SNP IDs) from the input **v**_*j*_’s into
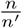 blocks, and process each block (of *n*′ dimensions) independently in the enclave memory. For a fixed *k*, the overall top-*k* dimensions (based on the standard *χ*^2^ statistic) can then be obtained by sorting the top-*k* dimensions of all blocks.

Unfortunately the above scheme does not allow the user to pose additional queries on processed data, e.g. identify the top *k*′-dimensions for *k*′ > *k* - either among the complete set of SNPs, or a user specified subset of SNPs. Here we describe an alternative solution implemented by `SkSESS`, which allows users to specify the value of *k* as well as a subset of SNPs of interest (among which the top-*k* dimensions need to be determined), even after all input data is processed. This solution maintains a “sketch” (or a summary) of the potentially important dimensions and allows the user to identify the top-*k* dimensions with respect to the *χ*^2^ values - with high probability.

`SkSES` processes the input vectors **v**_*j*_ in a single pass: for each (**v**_*j*_[*i*], sign(*j*)) it updates the appropriate entries of the sketch in an on-line fashion. The sketch basically maintains appropriate counts for *l* candidate dimensions (for a sufficiently large *l* > *k*) which include all top-*k* dimensions (with respect to the *χ*^2^ statistic) with high probability. Once the sketch is complete `SkSES` accesses the relevant vectors **v**_*j*_ in a second pass to filter out the false positives among these candidates and identify only the most significant *k* dimensions.

#### Definition 2.

Ranking SNPs with respect to *χ*^2^ statistic - with limited memory.

**Input:** A collection of *m* (sparse) vectors **v**_*j*_ with *n* dimensions where **v**_*j*_[*i*] ∈ {0, 1, 2}, each with a class label sign(*j*).

**Querying:** Again, there are two types of queries: return the top-*l* values among (1) rank(**Ĉ** [1: *n*])[1: *l*], or (2) rank(**Ĉ** [*i* ∈ *T* ⊂ [*n*]])[1: *l*], where **Ĉ** is an approximation to **C** (i.e. the *χ*^2^ values), such that the *k* largest values in **C** (respectively rank(**C**)[1: *k*] or rank(**C**[*i* ∈ *T*])[1: *k*]) are included.

Observe that

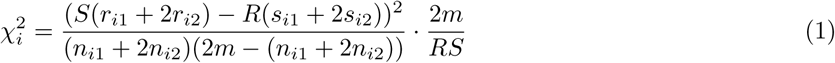

(which equals to
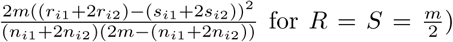 according to Definition 1. Let
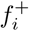 and
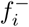 be the total allele counts for SNP *i* from case and control groups respectively, i.e.
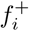 = *r*_*i*1_ + 2*r*_*i*2_,
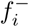 = *s*_*i*1_ + 2*s*_*i*2_; also let **F** be the vector where **F**[*i*] =
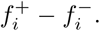 It follows that
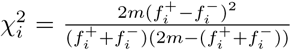 when *R* = *S*. Note that each (**v**_*j*_ [*i*], sign(*j*)) impacts
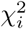 in a non-linear fashion. However, as will be shown below, we can rank the dimensions *i* with respect to the values
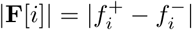 - as a proxy to
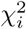 - since dimensions *i* with high
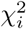 values (and thus are significant) can be demonstrated to have high |**F**[*i*]| values under some simple assumptions. (Note that when *R* ≠ *S* we need to rank the dimensions with respect to
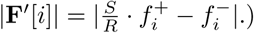

|**F**[*i*]| **as a proxy to
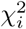.** Consider a GWAS setting where each dimension *i* ∈ [*n*] belongs to one of the two significance categories: *i* is a *low*-*significance* dimension, if the likelihood of a minor allele in a given individual is *ρ_i_*, independent of the case/control status of the individual; *i* is a *target* dimension if the likelihood of a minor allele in the case group, *ρ*_1*i*_, is significantly different from that in the control group, *ρ*_2*i*_.^3^ As demonstrated below, a target dimension *i* is likely to have a sufficiently large value for
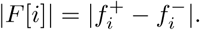 (Note that in case a target dimension *i* happens to exhibit a smaller |*F*[*i*]| value,
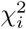 can not be high anyway.)

#### Lemma 1.

*For any target dimension i*, |**F**[*i*]| = Ω(|*ρ*_*i*1_ – *ρ*_*i*2_| · *m*) *with high probability.*

See the appendix for a proof.

#### Definition 2’.

Ranking SNPs with respect to *χ*^2^ statistic - with limited memory.

**Input:** A collection of 2*m* (sparse) *n* dimensional vectors **v**′_*j*_, where **v**′_*j*_ [*i*] {0, 1}, each with a class label sign(*j*).^4^

**Querying:** Return the top valued *l* dimensions with respect to (1) rank(|**F**|[1: *n*])[1: *l*], or (2) rank(|**F**|[*i* ∈ *T* ⊂ [*n*]])[1: *l*], where **F**: **F**[*i*] =
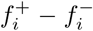 is an approximation to **C** (i.e. the *χ*^2^ values), such that the top valued *k* dimensions in **C** (within rank(**C**)[1: *k*] or rank(**C**[*i* ∈ *T*])[1: *k*]) are included.

### 2.7 The Sketching Data Structures for the SGX Enclave

In this section we describe two sketching data structures we implemented in the SGX enclave, which (i) generate unbiased estimate **F̂** to **F**, i.e. for each dimension *i*, 𝔼[**F̂**[*i*]] = **F**[*i*]; (ii) give a good approximation guarantee to each dimension *i* with a large |**F**[*i*]| =
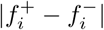 value so that those dimensions *i* with a high
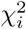 value will be among the candidate dimensions.

The two sketching data structures we consider are respectively based on the count-min sketch [18] and count sketch [17]. Let *w* and *d* denote the width and depth of the sketch (these are parameters whose values are determined later in Theorem 2); let *h*_1_, ⋯, *h_d_* be random pairwise independent hash functions to map each of the *n* dimensions to [*w*] (formally, *h_p_*(*i*): [*n*] → [*w*] for *p* ∈ [*d*]); similarly let *g*_1_, ⋯, *g_d_* be random pairwise independent hash functions that map each of the *n* dimensions to {–1, 1} (formally, *g_p_*(*i*): [*n*] → {–1, +1} for *p* ∈ [*d*] - this is used only in the count sketch). Both sketches consist of a *d* ∗ *w* matrix, where each entry represents a signed integer of *O*(log *m*) bits. Where the two sketches differ is in the update/query procedures (discussed below) and the performance guarantees they can provide. `SkSES` gives the users the ability to decide the sketch to be maintained in the SGX enclave.

Initially the entries in the *d* ∗ *w* matrix are all set to 0. To perform an on-line update in the form of (**v**_*j*_ [*i*], sign(*j*)), the count-min sketch data structure simply adds to the *h_p_*(*i*)-th column in the *p*-th row the value of sign(*j*) · **v**_*j*_ [*i*]. In contrast, the count sketch data structure adds to the same entry the value of *g_p_*(*i*) · sign(*j*) · **v**_*j*_ [*i*]. To obtain an unbiased query to each *i* ∈ [*n*], we apply a technique introduced in [41] as a special treatment for count-min sketch (interestingly the count sketch naturally gives unbiased estimates) – we maintain one extra counter *L* of *O*(log *m* + log *n*) bits to maintain the “data size”, to which we add sign(*j*) · **v**_*j*_ [*i*] at each update. Although each update step requires *d* increments or decrements (one in each row), they are trivially parallelizable since the operations to each row are independent. `SkSES` indeed employs a multi-threaded update mechanism, even though multi-threading in the SGX enclave comes with certain limitations, as discussed in the Results section.

After processing all updates, a query from count sketch to dimension *i* returns **F̂**[*i*] = median_*q*∈[*d*]_{*g_q_*(*i*) · *h_q_*(*i*)}; similarly, a query from count-min sketch to dimension *i* returns **F̂** [*i*] = median_*q*∈[*d*]_
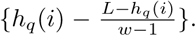 Before identifying the top-*k* significant SNPs, through processing the input data in a second pass, the user needs to specify the subset of SNP IDs of interest. Ideally, for *c* = Ω(min_*i*:target_{|*ρ*_*i*1_ – *ρ*_*i*2_|}), we should keep all the dimensions (SNP IDs) *i* with |**F̂**[*i*]| ≥ (*c* – *ϵ*)*m* to guarantee all candidate dimensions *i* with |**F**[*i*]| ≥ *c* · *m* are returned with high probability, which requires little extra space (in addition to the sketches) as analyzed in Appendix. In practice, `SkSES` simply maintains a min heap with at most *l* dimensions with the largest **F̂** values. Let min = min_*i*∈heap_{**F̂**[*i*]} be the smallest value in the current heap. To process a new dimension *i*′, if there are less than *l* dimensions in the heap we insert it into the heap; otherwise we replace *i*^∗^ = arg min_*i*∈heap_ **F̂**[*i*] when the query of *i*′ returns **F̂**[*i*′] > min.

`SkSES` allows a user to select the parameter *w* and *d*. The following theorem, however, gives the recommended parameters to obtain guaranteed accuracy performance.

#### Theorem 2.

*With high probability*, *(i) the query to count*-*min sketch returns all candidate dimensions by setting w* =
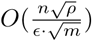 *and d* = *O*(log *n*)*; (ii) the query to count sketch returns all candidate dimensions by setting w* =
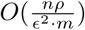 *and d* = *O*(log *n*), *where*
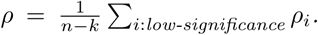 *The respective space usage of count*-*min and count sketch are*
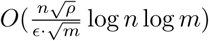 *and w* =
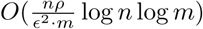 *bits*.

See the appendix for a proof.

## 3 Results

We tested our basic hashing schemes (see Section 2.5) and sketching data structures (see Section 2.7) on the iDASH-2017 (Track 2) [42] dataset, consisting of 1000 VCF files labeled “case” and another 1000 labeled “control” (i.e., *R* = *S* = 1000). The total size of the 2000 uncompressed VCF files were ~325G; overall they included *n* = 6.93 ∗ 10^7^ unique SNPs. We also used the 5.51 ∗ 10^6^ unique SNPs from chromosome 1 (the original benchmarking dataset for iDASH-2017 competition) in the 2000 VCF files for comparing the speed of hash tables with that of the sketching data structures. For convenience, the units (in particular, the parameters for sketches) in this section are denoted in powers of 2, i.e. 1*M* = 2^20^.

All of our experiments were run on a Linux server equipped with an SGX capable Intel Core i7-6700 (8 cores at 3.40GHz) processor with available physical memory of 8GB, although the Intel SGX enclave can only use 128MB due to the restrictions imposed by the architecture.^5^

### 3.1 Compression Efficiency

We applied our filtering and compression methods described in Section 2.3 first to the iDASH-2017 dataset for chromosome-1 and later to the whole genome. To simulate the scenario that each VCF file encodes a distinct patient whose information is private, we ran our filtering and compression algorithms on each file independently. As can be seen in Table 1, our filtering and compression approach results in roughly a 50-fold improvement over the plaintext in terms of data size.

**Table 1:** A comparison of our compression and filtering methods on the iDASH-2017 dataset. Top: compression ratios achieved on chromosome-1. Bottom: compression ratios achieved on the whole genome. An important distinction between our filtering/compression approach and that of gzip (tar.gz) is that we process each VCF file independently whereas we allowed gzip to process all VCF files jointly. In the very likely scenario that the VCF files originate from two or more institutions, they could not be compressed jointly. However, even though this comparison is unfair to our filtering/compression methods, we get an order of magnitude better compression results in comparison to those obtained by gzip.

| Dataset | Compression Method | Size (Bytes) | Ratio |
| --- | --- | --- | --- |
| iDASH2017-chr1 | None (Plaintext) | 28,975,640,907 Bytes | N/A |
| iDASH2017-chr1 | tar.gz (All Files Together) | 5,413,837,568 Bytes | 5.35 |
| iDASH2017-chr1 | Filtered + Zygoty Sorted (Each File Separately) | 2,321,993,336 Bytes | 12.48 |
| iDASH2017-chr1 | Filtered + Minimum Binary Representation | 574,648,245 Bytes | 50.42 |
| iDASH2017-all | None (Plaintext) | 349,709,350,603 Bytes | N/A |
| iDASH2017-all | tar.gz (All Files Together) | 68,909,020,026 Bytes | 5.08 |
| iDASH2017-all | Filtered + Zygoty Sorted (Each File Separately) | 29,589,341,724 Bytes | 11.82 |
| iDASH2017-all | Filtered + Minimum Binary Representation | 7,525,796,105 Bytes | 46.47 |

### 3.2 Runtime Performance

We evaluated the running time of our algorithms on the iDASH-2017 dataset (the data structure construction and query times were evaluated separately). On chromosome-1, our sketching algorithms outperform the standard hash table based methods. We need to mention here that this comparison under represents the running time advantage offered by the sketching based methods as the hash tables only report the top-10 SNPs with respect to the *χ*^2^ values, while the sketching schemes are capable of returning a much larger set of most significant SNPs (we set *l* = 1.3 ∗ 10^5^ here). On the whole genome, it is not possible to use the standard hash tables for dynamic queries due to memory restrictions, therefore we only report the running time results for our sketching data structures. As can be seen in Table 2, our sketching data structures handle the increase in the data size quite well with respect to both data structure construction and query times.

**Table 2:**
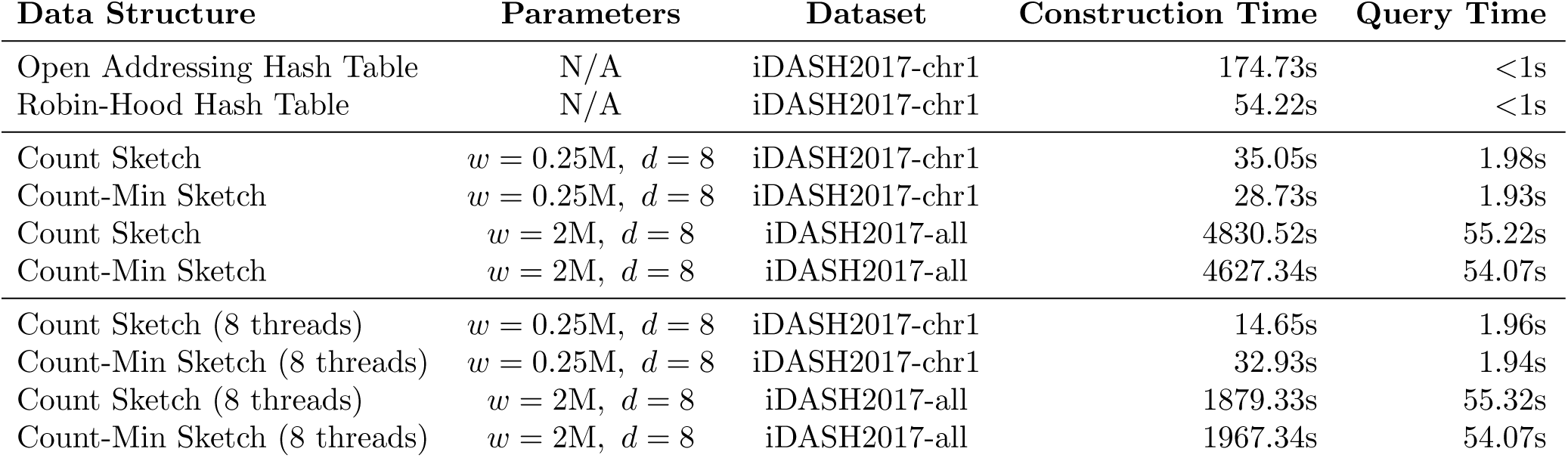
Running time comparison for all data structures on the iDASH-2017 dataset. Top: running times of the standard data structures; Middle: running times of the sketching data structures; Bottom: running times of the sketching data structures with 8 threads. Note that the standard data structures only report the top-10 SNPs with respect to the *χ*^2^ values whereas the sketching data structures report top-*l* estimates, where *l* can be as large as 1.3 ∗ 10^5^ (a smaller value of *l* leads to a better query time), thus their performance is significantly better than standard hashing approaches. Note that 1M = 2^20^ in the table.

As mentioned earlier, we also implemented and tested multi-threaded construction of the sketching data structures in SGX enclave. However, the multi-threading scheme for Intel SGX comes with the limitation that the number of enclave threads cannot exceed the number of logical processors, since an Intel SGX thread is directly mapped to a logical processor. For instance, consider a a multi-core system with 2 processors and each processor supporting 4 cores (this is akin to the system we used to obtain our performance figures). Then, as there is a one-to-one mapping between the enclave thread and the logical processor, the number of enclave threads would be limited to 8 for this system.

### 3.3 F as a proxy to C

As a first step in evaluating the performance of our sketching approach, we verified whether the top-*k* dimensions in rank(**C**) (i.e. the top *k* in **C**) were all included in the top-*l* dimensions in rank(|**F**|). We note here that **F** can be used as a proxy to **C** only if (i) each of the top *k* dimensions in **C** should have a sufficiently large |**F**| value; (ii) rank(**C**) and rank(|**F**|) should be well correlated.

As per Figure 1A, the minimum |**F**| value for any of the top 100 significant SNPs across all chromosomes is larger than 0.35*m* (*m* = 2000); very few SNPs outside of the top 100 have larger |**F**| values (i.e. lower rank). As *k* becomes larger, however, more SNPs need to be maintained so that the true top *k* could be returned. Alternatively, if we allow (in the answer to queries with larger *k*) a small fraction of “false negative” SNPs (not not returned by `SkSES`) the number of “false positive” SNPs are significantly reduced; see Figure 1B demonstrating this tradeoff.

**Figure 1:**
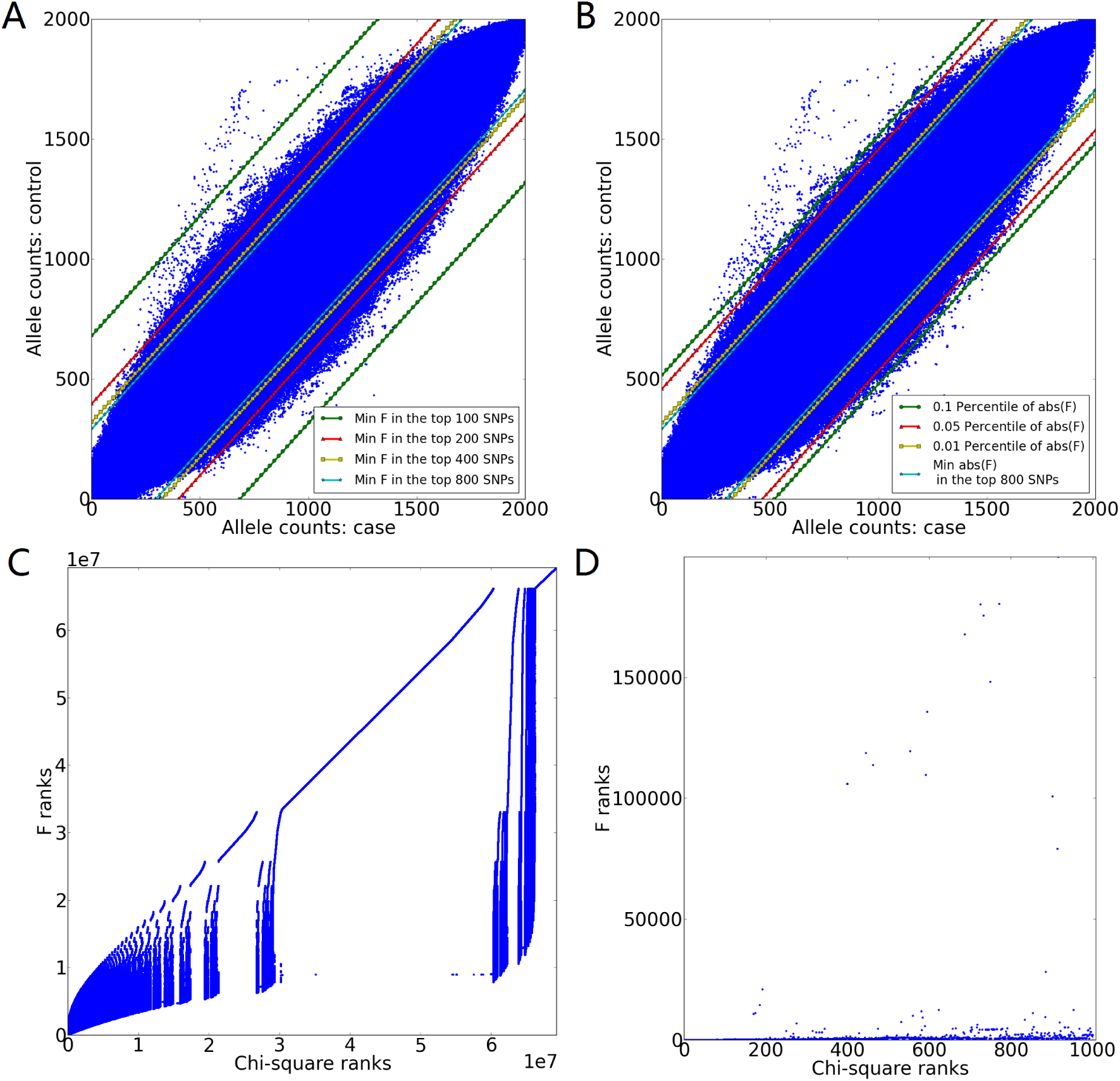
Correlation between the rank of a SNP with respect to the *χ*^2^ statistic (i.e. rank(**C**)) and the rank with respect to |**F**| (i.e. rank(|**F**|)) across all unique SNPs from the whole iDASH2017 dataset. (A) The minimum |**F**| value given by the top *k* significant SNPs according to *χ*^2^ statistic, for varying *k*’s. (B) The minimum |**F**| value given by the top *k* significant SNPs according to *χ*^2^ statistic, for *k* = 800, with varying “false negative” (i.e. the SNPs are among the true top *k* ones but not reported by `SkSES`) rates. In both (A) and (B), a blue point represents a particular SNP *i* with allele count
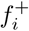 in the case group and allele count
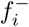 in the control group. (C) Correlation between the rank(**C**) and rank(|**F**|), where x-axis gives the rank of a particular SNP *i* in **C** and y-axis gives its rank in |**F**|. (D) Correlation between the rank(**C**) and rank(|**F**|) for the top 1000 SNPs according to **C** (the *χ*^2^ statistic).

Figure 1C and Figure 1D demonstrate the correlation between rank(**C**) and rank(|**F**|). As can be seen in Figure 1C, rank(|**F**|) provides an approximation of rank(**C**) without false negatives, as there are no SNPs with significantly higher ranks with respect to |**F**| than their rankis with respect to **C**. Orthogonally, Figure 1D demonstrates a (small) constant *α* such that if we keep the first *αk* SNPs in rank(|**F**|), the true top *k* SNPs in **C** are almost guaranteed to be included (the number of “false negative” SNPs depends on *α*), for reasonably small values of *k*. As a result, the total number of SNPs needed to be kept to include all top-*k* SNPs, is also bounded, provided that **F** can be queried accurately. We discuss the overall “false negative” rate obtained by `SkSES` due to the approximation in **F** in the next section.

### 3.4 Accuracy of Sketches

We evaluated the percentage of the true top *k* SNPs (w.r.t the *χ*^2^ statistic) as a function of *l*, the number of SNPs returned by the sketches (denoted as “accuracy”) for both chromosome 1 and the whole genome for varying sketch sizes. As can be seen in Figure 2, a wider and deeper sketch results in increased accuracy, although a narrower and more shallow sketch may offer a better memory utilization and result in faster running time. Due to the restrictions imposed by SGX enclaves (explained in Section 3.2), selecting a depth greater than the maximum number of threads supported by the enclave could result in significant decline in performance, therefore we used *d* = 8, setting the sketch depth to the maximum number of threads supported by an SGX enclave. Even under such conditions, the queries to sketches still offer reasonably high accuracy. Interestingly, our sketching data structures always return the top 100 SNPs w.r.t the *χ*^2^ statistic with 100% accuracy further supporting the statistical significance of the top 100 SNPs in iDASH2017 dataset, both from chromosome 1 and the whole genome. For larger values of *k*, the accuracy degrades slower for larger sketches, leading to a trade-off between speed and accuracy.

**Figure 2:**
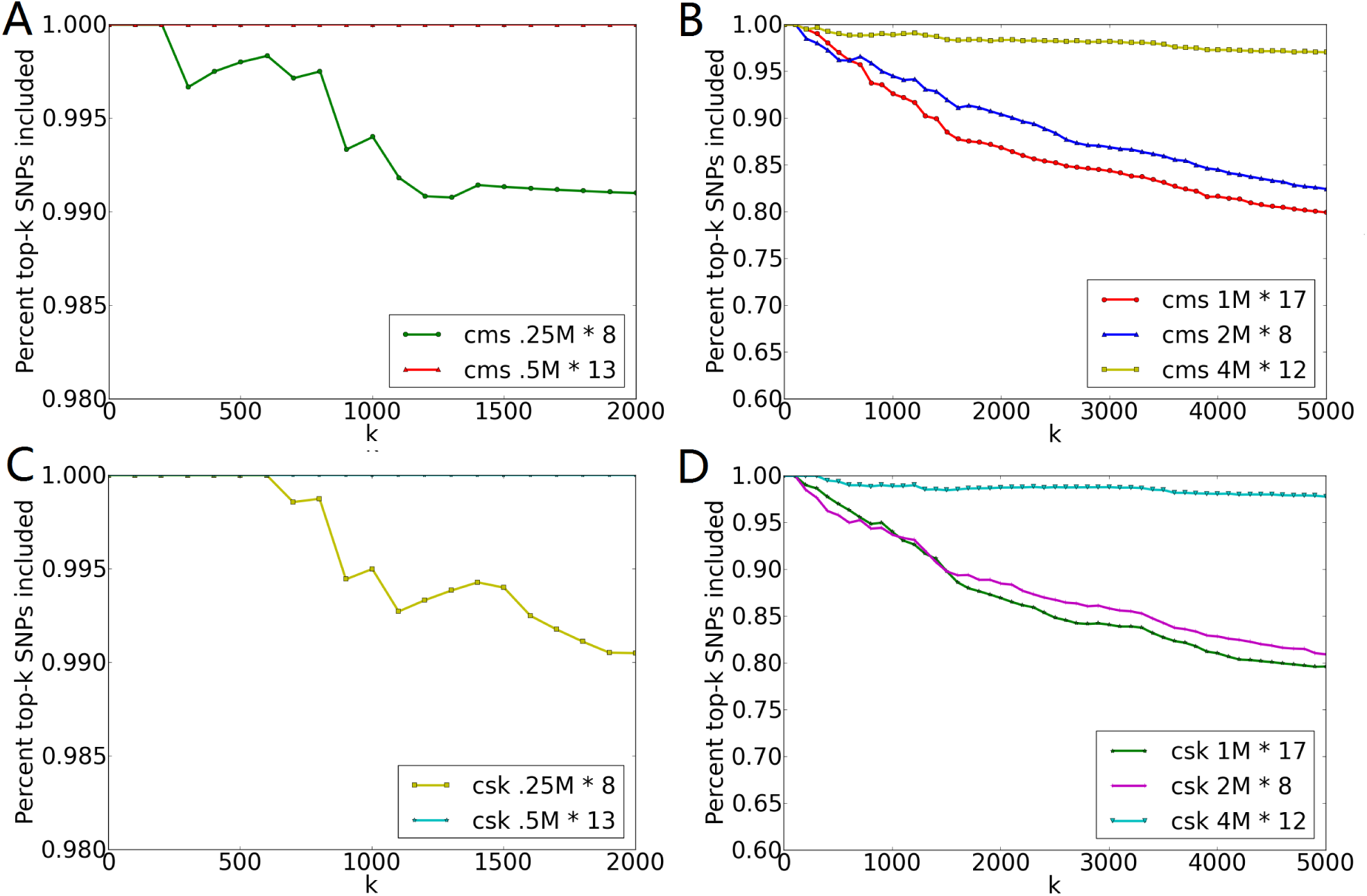
The percent of the true top *k* significant SNPs (according to *χ*^2^ statistic) included in the query of top *l* SNPs returned by `SkSES` (i.e. “accuracy”), as a function of *k*. The queries were either from count-min sketch (A, B), or from count sketch (C, D), for chromosome 1 only (A, C) and the whole genome (B, D) in the iDASH2017 dataset, with the same parameter *l* = 1.3 ∗ 10^5^. Other parameters for the sketches include the width *w* and depth *d*, as shown in each subplot, where 1M = 2^20^. The parameters were chosen in a way that either the speed for sketch updates and queries is fast enough, or the accuracy for querying with small *k*’s is plausible, while the sketches can still fit into the memory provided by secure enclaves.

## 4 Conclusion

In this paper we describe `SkSES` a framework for the Intel SGX architecture to perform secure and privacy preserving GWAS in an untrusted cloud environment through the use of modified sketching data structures. In order to facilitate efficient use of the limited memory available within the SGX enclave, `SkSES` employs simple filtering and compression schemes that offer an order of magnitude improvement over plaintext representations. Our sketching algorithms offer better performance compared to standard hash table based approaches on chromosome-1 with near hundred percent accuracy. They also perform well on the whole genome - where standard hash table based methods can not support on-line querying and thus are infeasible.

## Appendix A. Proofs

Recall that there are *m* input vectors **v**_*j*_ ∈ {0, 1}^*n*^ (*j* ∈ [*m*]) labeled with “case”, and *m* labeled with “control” (*j* ∈ [2*m*] – [*m*]). Each vector is of dimension *n*, in which there are at most *k* target dimensions and at least *n* – *k* low-significance dimensions. Let *I* ⊂ [*n*] and *T* ⊂ [*n*] be the set of low-significance dimensions and target dimensions, respectively. Furthermore, let *V*_*j*,*i*_ be i.i.d. random variables indicating the value of **v**_*j*_[*i*]; *F_i_* = ∑_*j*∈[2*m*]_ *V*_*j*,*i*_. Then for *i* ∈ *I*, Pr[*V*_*j*,*i*_ = 1] = *ρ_i_*; and for *i* ∈ *T*, Pr[*V*_*j*,*i*_ = 1] = 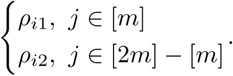 Finally, we let
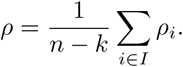

### Proof of Lemma 1

#### Proof.

For *i* ∈ *T*, 𝔼[*F_i_*] = (*ρ*_*i*1_ – *ρ*_*i*2_)*m*. It follows from Chernoff bound that
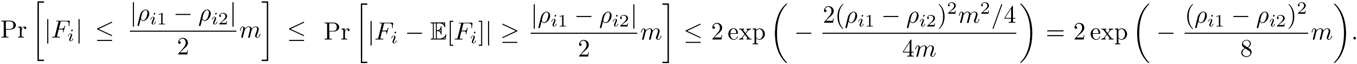 As *ρ*_*i*1_ and *ρ*_*i*2_ are constants depending on the input, the Lemma holds for sufficiently large *m*.□

### Lemma 3.

*For a low*-*significance dimension i*,
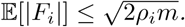

### Proof.

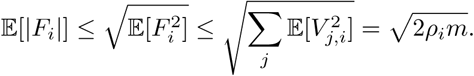□

### Lemma 4.

*With high probability*, *(1)* ∥**F**_–*k*_∥_1_ = *O*(*n* ·
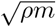)*; (2)* ∥**F**_–*k*_∥_2_ = *O*(*n* · *ρm*).

### Proof.

First notice that
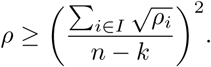 By Lemma 3,
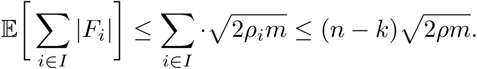 Thus, applying Chernoff bound, we have
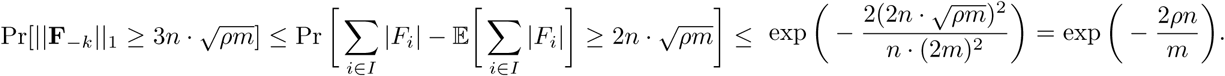 Typically *n* >> *m*, whereas *ρ* is a constant depending on the input **v**_*j*_’s, making the above “failure” probability sufficiently small. The proof to (2) follows in a similar fashion.□

### Theorem 5.

*([41*, *17]) (1) Both sketches return* **F̂**, *such that* 𝔼[**F̂**[*i*]] = **F**[*i*] *for each i* ∈ [*n*]*; (2) By setting d* = Θ(log *n*) *and w* = Ω(*k*), *we have*
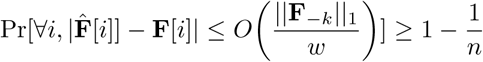 *for count*-*min sketch*, *and*
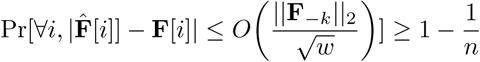 *for count sketch*, *where* **F**_–*k*_ *denote the vector* **F** *after zeroing out the k dimensions with the largest absolute values*.

### Proof of Theorem 2

#### Proof.

The space required by count-min sketch and count sketch giving *ϵm* error bound follows directly from Lemma 4 and Theorem 5, i.e., for count-min sketch,
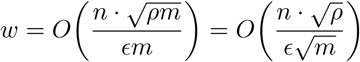 (and *d* = Θ(log *n*)), while for count-min sketch
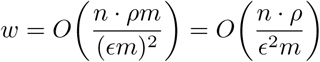 (and *d* = Θ(log *n*)). On the other hand, for a low-significance dimension *i* ∈ *I*, the probability that it is returned in a query is sufficiently small: Pr[|*F_i_*| ≥ (*c* – *ϵ*)*m*] ≤
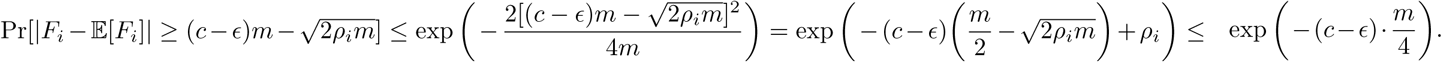 Therefore, compared to the space needed for maintaining the sketches, the space needed for keeping low-significance dimensions in addition to the *k* target dimensions is negligible, if we need to maintain all dimensions *i* with |**F̂**[*i*]| ≥ (*c* – *E*)*m*.

## Footnotes

1 `SkSES` identifies the most significant SNPs based on the *χ*^2^ statistic, the most commonly used in GWAS, but it can easily be modified to accommodate other statistics. However, since GWAS aim to find biologically meaningful associations between genomic variants and specific diseases, one may need to go beyond identifying the most significant SNPs - since biologically meaningful mutations are many times inherited together with variants that have no functional significance. For addressing this “population stratification” problem, the input data can be preprocessed (e.g. via Eigenstrat [39]) to reduce the impact of variational patterns specific to distinct human populations. We assume here that the input data is already preprocessed based on variational patterns observed in the general human population.

2 The current effective memory offered by the SGX enclave is 128MB.

3 In this setting the likelihood of a minor allele is assumed to be independent across distinct dimensions. Although this assumption simplifies our analysis, it does not hold in practice due to population stratification - however the impact of population stratification can be reduced through a number of available preprocessing methods as mentioned earlier.

4 For simplifying the problem, we assume that each vector **v**_*j*_ from the original input definition is replaced with two vectors **v**_*j*1_ and **v**_*j*2_ such that **v**_*j*1_ + **v**_*j*2_ = **v**_*j*_ w.l.o.g.

5 For Windows data sizes exceeding 128MB will cause the enclave to crash, whereas Linux supports paging. However crashes can still occur and paging is several orders of magnitude slower than regular memory access, and thus should be avoided.

